# Mouse conventional dendritic cells can be universally classified based on the mutually exclusive expression of XCR1 and SIRPα

**DOI:** 10.1101/012567

**Authors:** Stephanie Gurka, Martina Becker, Evelyn Hartung, Richard A. Kroczek

**Author notes:** contributed equally to this work (Stephanie Gurka, Martina Becker, Evelyn Hartung). Correspondence: Richard Kroczek, M.D. Molecular Immunology Robert Koch-Institute Nordufer 20, 13353 Berlin, Germany.

## Abstract

Since the identification of mouse dendritic cells (DC) in the early 70s, all attempts to consistently classify the identified functional DC subpopulations according to their surface molecule expression failed. In the absence of DC lineage markers, a great variety of non-congruent surface molecules were used instead. Recent advances in the understanding of the involvement of transcription factors in the differentiation of DC subpopulations, together with the identification of a lineage marker for cross-presenting DC, have now allowed to establish a consistent and unified DC classification in the mouse. We demonstrate in the present article that all conventional DC in the mouse can be universally subdivided into either XCR1^+^ cross-presenting DC or SIRPα^+^ DC, irrespective of their activation status. This advancement will greatly facilitate future work on the biology of mouse DC. We discuss this new classification in view of current DC classification systems in the mouse and the human.

## Overview

Dendritic cells (DC) were discovered by Steinman et al. already in the early 70s (Steinman and Cohn, 1973). Nevertheless, it was until recently difficult to unequivocally distinguish them from other related cell types such monocytes or macrophages. As a result, a combination of several markers had to be used to define DC in flow cytometry and histology. For practical purposes, mouse conventional DC today are identified in flow cytometry as cells which express the integrin CD11c and high levels of MHC II, but lack expression of T-, B-, and plasmacytoid DC lineage markers, and also molecules characteristic for monocytes and macrophages.

In the absence of (sub-) lineage markers, also DC subpopulations could not easily be defined using surface molecules. This has led over time to the use of a great variety of surface markers distinguishing supposedly functionally distinct DC subpopulations and made it difficult to directly compare results between laboratories. Further complexity was brought about by the observation that DC with an apparently similar function had different phenotypes in lymphoid tissues versus peripheral organs. As a consequence, the division of DC into subpopulations remained somewhat arbitrary, making the experimental results, including gene expression studies, less informative. Being central to the understanding of DC biology, the question how mouse conventional DC should be divided into subsets remained a matter of intensive scientific debate to this very day.

Recently, we identified the first molecule restricted in its expression to mouse DC (Dorner et al., 2009). Based on a variety of experimental systems, we then could demonstrate that the chemokine receptor XCR1 is a lineage marker for cross-presenting DC (Bachem et al., 2012), a DC subpopulation playing an important role in the induction of CD8^+^ T cell cytotoxicity (see below). The use of an antibody directed to XCR1 allowed for the first time the unequivocal identification and thus a precise phenotyping of cross-presenting DC in various body compartments of the mouse (Becker et al., 2014b; Cerovic et al., 2014, and Figures 1 and 2).

**Figure 1.**
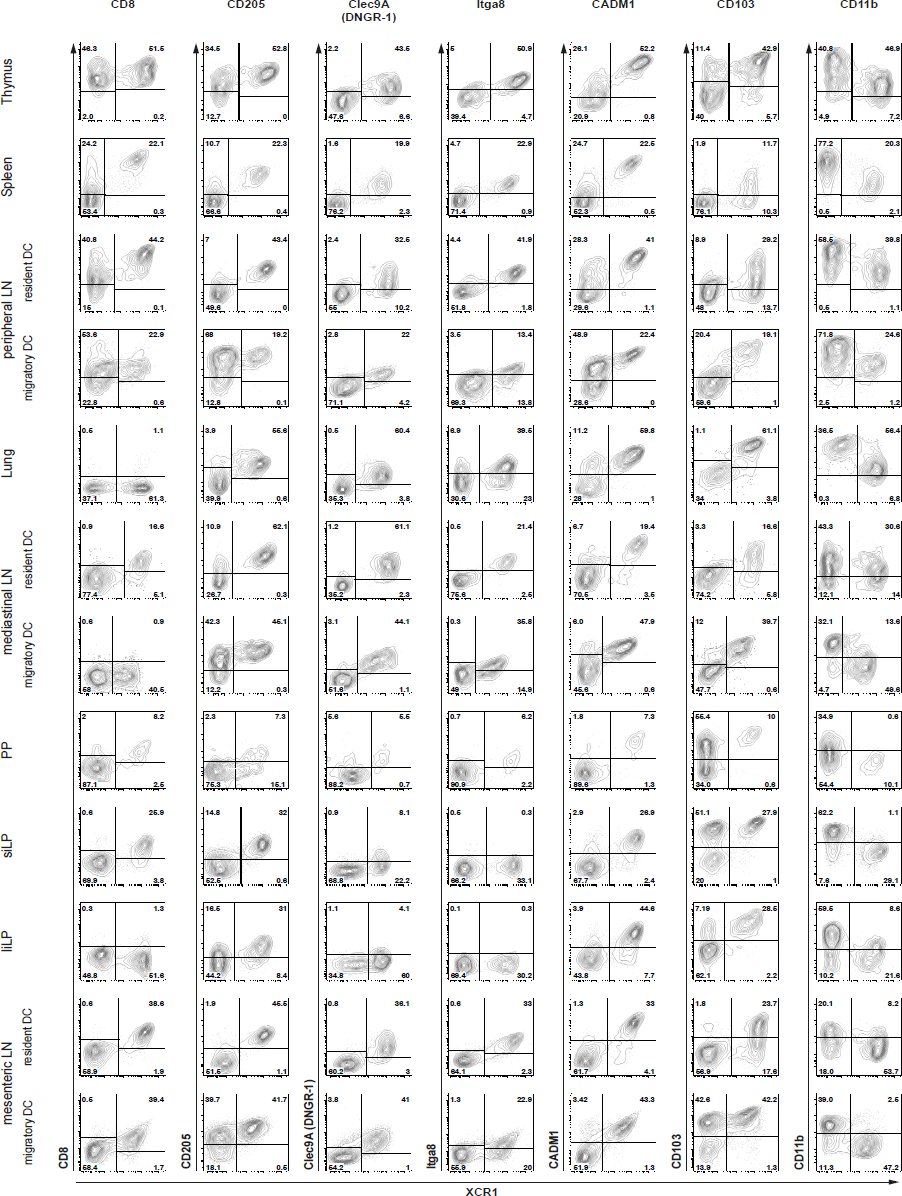
Correlation of XCR1 expression with different DC surface molecules at steady state. Cells from different tissues of C57BL/6 mice were isolated after enzymatic digestion (except for the spleen), and DC from lamina propria (LP), Peyer’s patches (PP), and mesenteric LN were additionally enriched by density gradient centrifugation as described before (Bachem et al., 2012; Becker et al., 2014b). Cells from brachial, axillary, and inguinal LN were pooled for the study of peripheral LN DC. For flow cytometric analysis of XCR1 and SIRPα expression on DC, gates were set on live CD90^−^ CD19^−^ CD317^−^ CD11c^+^ MHC II^+^ cells for thymus, spleen, peripheral and mediastinal LN, and on live CD45^+^ CD3^−^ B220^−^ F4/80^−^ CD11c^+^ MHC II^+^ cells for lung, Peyer’s patches, LP, and mesenteric LN. DC from LN were separated based on their MHC II expression levels into resident (MHC II^int^) and migratory (MHC II^high^) populations. liLP, large intestinal lamina propria; siLP, small intestinal lamina propria.

**Figure 2.**
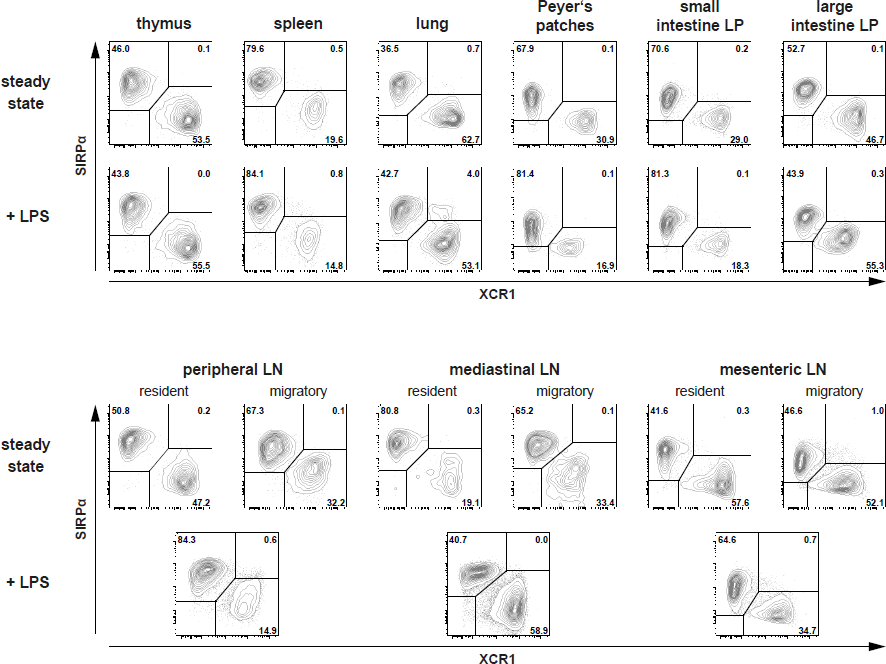
Expression of XCR1 and SIRPα on DC in various tissues at steady state and in inflammation. Cells from different tissues of C57BL/6 mice (either untreated or 14 h after injection of 10 μg LPS i.v.) were isolated after enzymatic digestion (except for the spleen), and DC from lamina propria (LP), Peyer's patches, and mesenteric LN were additionally enriched by density gradient centrifugation as described before (Bachem et al., 2012; Becker et al., 2014b). Cells from brachial, axillary, and inguinal LN were pooled for the study of peripheral LN DC. For flow cytometric analysis of XCR1 and SIRPα expression on DC, gates were set on live CD90^−^ CD19^−^ CD317^−^ CD11c^+^ MHC II^+^ cells for thymus, spleen, peripheral and mediastinal LN, and on live CD45^+^ CD3^−^ B220^−^ F4/80^−^ CD11c^+^ MHC II^+^ cells for lung, Peyer’s patches, LP, and mesenteric LN. DC from steady-state LN were separated based on their MHC II expression levels into resident (MHC II^int^) and migratory (MHC II^high^) populations.

With the ability to define the cross-presenting DC population, it became possible to ask the question whether the remaining DC could also be defined by their surface characteristics. The use of an extended panel of antibodies directed to DC surface molecules indicated that all XCR1^−^ DC were characterized by expression of SIRPα/CD172a (Becker et al., 2014b). Based on these results, together with the data presented here, we now propose a new classification based on the expression XCR1 and SIRPα, which can be used to define DC subpopulations in all lymphoid and non-lymphoid compartments of the mouse.

This article describes the various steps which have led to the establishment of this new DC classification system, and discusses the implications for the understanding of human DC subpopulations. Gene expression profiles and functional aspects of DC subsets (recently reviewed by Hashimoto et al., 2011; Haniffa et al., 2013; Merad et al., 2013; Mildner and Jung, 2014) are taken into account only as far as they contribute to the classification of mouse DC. Monocyte-derived inflammatory DC and plasmacytoid DC are not considered here, since they represent different cell lineages.

## Historical DC classification systems

Work of numerous groups has established that at least two major conventional DC populations exist in the mouse. In the late 90s, a subset of DC was identified in lymphoid tissues which expressed the CD8α homodimer on the cell surface (and lacked CD8β and CD11b), and these DC were hence termed “CD8^+^ DC” (Shortman and Heath, 2010). Major steps forward in the understanding of DC biology was the demonstration that CD8^+^ DC excel in antigen “cross-presentation”, in which antigen is not presented in the context of MHC II to CD4^+^ T cells, but instead shunted into the MHC I pathway and presented to CD8^+^ T cells (Kurts et al., 1996, den Haan et al., 2000; Pooley et al., 2001). Further studies demonstrated that mouse CD8^+^ DC are specialized in the uptake and proteolytic processing of stressed cells and the subsequent presentation of the derived peptides to CD8^+^ T cells (Iyoda et al., 2002; Schulz and Reis e Sousa, 2002; Schnorrer et al., 2006). In general terms, antigen cross-presentation allows an efficient induction of CD8 T cell cytotoxicity to antigens originating from cell-invading pathogens or mutated (cancer) cells.

After identification of CD8α as a relevant subset marker on around 20% of DC in the spleen, the remaining splenic DC were classified as CD4^+^ DC (60%) and double-negative DC (DN DC, 20%) (Vremec et al., 2000). When Edwards et al. (Edwards et al., 2003) performed microarray gene expression profiling of splenic DC populations in the mouse, they confirmed the differing nature of CD8^+^ DC, but noticed only relatively minor differences in the gene expression profiles of CD4^+^ and DN DC, and thus concluded that these two populations are phylogenetically related. Later work based on targeting of antigen directly to splenic DC subsets has confirmed the superior capacity of CD8^+^ (DEC-205^+^) DC for antigen cross-presentation and at the same time demonstrated a higher efficiency of CD8^−^ (33D1/DCIR^+^) DC in the presentation of antigen to CD4^+^ T cells (Dudziak et al., 2007).

While the phenotypical and functional classification of splenic DC made substantial progress, the understanding of DC subpopulations in peripheral lymphoid tissues and organs lagged behind. CD8α was detectable on resident DC in all lymphoid organs, but absent on DC in certain peripheral tissues and on DC migrating from the periphery to lymph nodes (LN) (Sung et al., 2006; del Rio et al., 2007). Since CD8α was less useful in the periphery, a different classification system was established by subdividing DC into CD103^+^ and CD103^−^ populations (Annacker et al., 2005; del Rio et al., 2007; Jaensson et al., 2008; Bedoui et al., 2009). Antigen cross-presentation was shown to be restricted to CD103^+^ DC residing in the lung, the intestine, and skin-draining LN (del Rio et al., 2007; Jaensson et al., 2008; Bedoui et al., 2009), suggesting a functional relationship to the CD8^+^ splenic DC. However, it soon was recognized that the CD103^+^ DC population is not homogenous and therefore the exact relationship between peripheral DC and lymphoid-resident DC remained unresolved.

## Involvement of transcription factors in the differentiation of DC

A major step forward was brought about by work on the involvement of transcription factors (TF) in the differentiation of DC. A series of studies demonstrated that development of CD8^+^ splenic DC and their peripheral counterparts critically depend on the TF IRF-8 (also designated ICSBP), Id2, and Batf3 (Schiavoni et al., 2002; Aliberti et al., 2003; Hacker et al., 2003; Hildner et al., 2008; Ginhoux et al., 2009; Edelson et al., 2010). The most informative turned out to be the Batf3-KO mouse, where only the splenic CD8^+^ DC, the lung and dermal CD103^+^ DC, and the intestinal CD103^+^ CD11b^−^ DC were absent and thus could be identified as developmentally related (only later it became apparent that other small DC populations with a differing phenotype were also Batf3-dependent, see below). At the same time, this animal model showed clear deficits in antigen cross-presentation (Hildner et al., 2008; Edelson et al., 2010). Together, this work strongly indicated that the Batf3-dependent DC were the cross-presenting DC lineage in the mouse.

## Identification of a lineage marker for cross-presenting DC

When searching for the function of XCL1, a chemokine secreted by activated CD8^+^ T cells and NK cells (Müller et al., 1995; Dorner et al., 2002), we found that the corresponding receptor XCR1 is exclusively expressed by a subset of DC. This observation represented the first instance of a surface molecule being restricted to conventional DC in the mouse. Analyzing a XCR1-lacZ-reporter mouse using flow cytometry, a system which provides limited signal resolution and high background in some extra-splenic tissues, we found XCR1 to be expressed by 70-90% of CD8^+^ DC and by up to 8% of DN splenic DC. Histological analyses further indicated that other lymphoid tissues and peripheral organs contained XCR1^+^ DC (Dorner et al., 2009). Using the same XCR1-reporter mice and extending the flow cytometry studies to DC in lymph nodes and several organs, Crozat et al. (Crozat et al., 2011) found XCR1 signals essentially limited to CD103^+^ CD11b^−^ DC, allowing them to postulate that expression of XCR1 defines mouse lymphoid-tissue resident and migratory DC of the “CD8α-type”.

Further understanding of XCR1-expresssing DC became possible with the development of a mAb specific for murine (and rat) XCR1, which offered high-resolution flow cytometry and also allowed sorting of XCR1^+^ DC for functional assays. These studies (Bachem et al., 2012) confirmed earlier findings with the lacZ-reporter system in the spleen (Dorner et al., 2009) that expression of XCR1 and CD8 overlap, but are not congruent. In the lung, the intestine, in skin-draining and mesenteric LN, CD103^+^ CD11b^−^DC were found to be essentially XCR1^+^. However, additional XCR1^+^ DC populations could also be identified there and these were negative for CD103 or positive for CD11b (Bachem et al., 2012; Becker et al., 2014b). Thus, the expression pattern of XCR1 differed from the “CD8^+^” and “CD103^+^ CD11b^−^” DC phenotypes associated with antigen cross-presentation in the past.

In experiments directly aimed to define the correlation between XCR1 expression and Batf3-dependence of DC, it became apparent that all XCR1^+^ DC (irrespective of their CD8, CD103, or CD11b expression status), were absent in Batf3-KO mice (Bachem et al., 2012; Becker et al., 2014b). Congruent with this observation, the 20% of splenic CD8^+^ DC, which are negative for XCR1, were preserved in Batf3-KO animals (Bachem et al., 2012); these particular CD8^+^ XCR1^−^ DC apparently represent a distinct DC population with a very different gene expression profile (Bar-On et al., 2010). Together, the studies demonstrated for DC in all tissues an excellent correlation between XCR1 surface expression and dependence on Batf3.

These correlation studies were very striking, but did not deliver direct information on the functional role of XCR1^+^ versus XCR1^−^ DC. Only when functional assays using soluble and cell-associated antigen were performed with splenic (Bachem et al., 2012) and intestinal DC (Becker et al., 2014b; Cerovic et al., 2014), it became apparent that antigen cross-presentation is restricted to XCR1^+^ DC, irrespective of their additional phenotype. Conversely, the CD8^+^ DC negative for XCR1 were found to be incapable of antigen cross-presentation (Bachem et al., 2012).

The perfect correlation between expression of XCR1, Batf3-dependence, and the ability to cross-present (cell-associated) antigen in various organ systems (Bachem et al., 2012; Becker et al., 2014b; Cerovic et al., 2014) allow to conclude that XCR1 expression generally demarcates the Batf3-dependent cross-presenting DC, as postulated (Bachem et al., 2012). Thus, XCR1 can be regarded as the lineage marker for cross-presenting DC in the mouse.

## Alternative markers for cross-presenting DC?

Could other surface molecules be used to demarcate these mouse cross-presenting DC? The shortcomings of CD8α as a DC-marker were discussed above. CD205, an endocytic recognition receptor for dead cells (Shrimpton et al., 2009) was also often used to define cross-presenting DC in the past. It does correlate quite well with XCR1 in most tissues, however, in migratory DC present in skin-draining and mesenteric LN or in the thymus, the correlation is poor, since additional DC express CD205 (Becker et al., 2014b, and Figure 1). Clec9A/DNGR-1, in the mouse expressed on DC and on plasmacytoid DC only (Caminschi et al., 2008; Sancho et al., 2008), is a receptor which has been shown to optimize processing of dead cells (Sancho et al., 2008). On conventional DC, Clec9A/DNGR-1 is perfectly correlated with the surface expression of XCR1 (Bachem et al., 2012; Becker et al., 2014b), but often is detectable at very low levels only (Becker et al., 2014b, and Figure 1), precluding its isolated use for the demarcation of cross-presenting DC. Tissue expression of the integrin Itga8 appears to be rather restricted to DC and some stromal cells (Immgen database). If Itga8 can be detected on DC, its low-level expression appears to correlate very well with XCR1, but in some instances (e.g. large and small intestinal lamina propria DC) no signal on XCR1^+^ DC could be obtained (Figure 1). Finally, the cell adhesion molecule CADM1 appears to be highly but not fully correlated with XCR1, since cells expressing low levels of CADM1 but no XCR1 can be detected in various organs (Figure 1). Thus, mouse cross-presenting DC can be best delineated by expression of XCR1.

## All DC can be classified into XCR1^+^ versus SIRPα^+^ DC irrespective of their activation state

Is there a surface molecule which would define the remaining, the XCR1^−^ DC population? To examine this question, we reanalyzed all of our correlation studies which included a large panel of antibodies directed to DC surface molecules, among others CD11b, CD172a/SIRPα, DCIR2, CD207, and a CX3CR1-reporter mouse. In all of our analyses, the only molecule which showed a consistent and full anti-correlation with XCR1 was CD172a/SIRPα, indicating that this surface molecule could be used to positively demarcate XCR1^−^ DC (Bachem et al., 2012; Becker et al., 2014b). Based on these studies, we have proposed that XCR1 and SIRPα can be used to classify intestinal DC and possibly all DC in the mouse (Becker et al., 2014a; Becker et al., 2014b).

In order to test the general applicability of this new classification system and to make the XCR1 expression studies directly comparable, we have now isolated DC from a greater variety of lymphoid and non-lymphoid organs and analyzed them in parallel. As can be seen in Figure 2, XCR1 and SIRPα were found to be clearly anti-correlated in all organs tested. At the same time, all DC present in these organs could be assigned to either population. Thus, the anti-correlation between XCR1 and SIRPα can now be demonstrated in a great variety of tissues.

All published data on the anti-correlation of XCR1 and SIRPα have been obtained in steady-state animals only. It was therefore important to test DC also under inflammatory conditions, when many DC surface molecules are up or down regulated. To this end, animals were injected with 10 μg LPS i.v. and DC examined 14 h later. Under these conditions, SIRPα expression remained rather stable, while XCR1 was slightly down regulated in some organs, however without compromising the discrimination of XCR1^+^ DC from SIRPα^+^ DC (Figure 2). Thus, the subdivision of conventional DC based on the expression of XCR1 and SIRPα appears to be universally applicable. With the commercial availability of an antibody directed to mouse (and rat) XCR1 (clone ZET), this classification can now be generally tested under all possible conditions.

SIRPα, also abundantly expressed on macrophages, neutrophils and some non-lymphoid tissues, has been implicated in the control of cell phagocytosis (Matozaki et al., 2009; Nuvolone et al., 2013). Cells expressing CD47, the ligand for SIRPα, appear to be protected from engulfment by phagocytic cells (Matozaki et al., 2009). It is thus intriguing to note that both CD205 and Clec9A/DNGR-1 on XCR1^+^ DC regulate phagocytosis in a positive, and SIRPα on XCR1^−^ DC in a negative fashion. This functional feature may possibly contribute to the division of labor between the XCR1^+^ and SIRPα^+^ DC populations.

All available data (gene expression profile, toll-like receptor expression pattern in particular), indicate that XCR1^+^ DC are a homogenous DC lineage. Does this mean that all XCR1^+^ DC function in the same way? This may not be the case. It is of interest in this context that only CD8^+^ (i.e. XCR1^+^) splenic DC expressing CD103 were capable to take up cells and to cross-present their antigen to CD8^+^ T cells in a previous study (Qiu et al., 2009). Only approximately 50–60% of splenic CD8^+^/XCR1^+^DC express CD103 under non-inflammatory conditions (Qiu et al., 2009; Bachem et al., 2012), and the situation appears similar in resident mesenteric LN (Pribila et al., 2004; Becker et al., 2014b). It is therefore quite possible that XCR1^+^ DC are functionally heterogenous in terms of cross-presentation, depending on their activation state, their anatomical positioning, and the upregulation of other molecules in reaction to environmental cues.

## Are SIRPα^+^ DC homogenous in their ontogeny and function?

This question is largely unresolved at present. Edwards et al. (Edwards et al., 2003) showed that CD4^+^ DC and DN DC, which now would be grouped together as SIRPα^+^ DC, have a highly similar gene expression profile, suggesting one uniform population. Nevertheless, the authors found some genes to be quite specifically expressed in either population. Other reports also point to a possible ontogenic or functional subdivision of SIRPα^+^ DC. For example, only a small fraction of SIRPα^+^ DC in the spleen express CD8 and differ in their expression profile from the remaining DC populations (Bar-On et al., 2010; Bachem et al., 2012). Mice with a CD11c-cre driven deletion of the TF Notch2 showed a rather specific ablation of CD103^+^ CD11b^+^ DC in the intestinal system (representing an ablation of approximately 50% of SIRPα^+^ DC), and these DC were identified as an obligate source of IL-23 in the defense to *C. rodentium* (Satpathy et al., 2013). In another study, depletion of only CD301b^+^ DC in the CD103^−^ (and thus most likely SIRPα^+^) fraction of dermal DC resulted in a severe impairment of skin Th2 immunity (Kumamoto et al., 2013). More studies will be required to make the results comparable and to answer the question on the heterogeneity of SIRPα^+^ DC.

## Can the subdivision of DC into XCR1^+^ and SIRPα^+^ DC be also applied in the human?

In humans, in vivo experiments on the function of DC subsets are not possible, access to DC in various compartments is limited, and the frequency of DC in the blood is very low. As a result, data on human DC are rather scarce. At the gene expression level, it was well established that mouse CD8^+^ DC correspond to human CD141^+^ (BDCA3^+^) DC, and mouse CD11b^+^ DC to human CD1c^+^ (BDCA1^+^) DC (Robbins et al., 2008), the two identified human DC populations (Ju et al., 2010). Support for this correlation came from functional studies which demonstrated a superior capacity of CD141^+^ DC for cross-presentation (Bachem et al., 2010; Crozat et al., 2010; Jongbloed et al., 2010). However, gene expression studies rely on previous sorting of DC populations and thus depend on the use of “correct” surface markers. Regarding these surface markers, however, human DC somewhat differ from mouse DC. XCR1 is exclusively expressed on CD141^+^DC, but not on all of them. Analyses of DC obtained from peripheral blood, thymus, and spleen demonstrated that an average of 80% of CD141^+^DC express XCR1 (own unpublished data). At the same time, all of the CD141^+^ DC express Clec9A/DNGR-1, which in the human appears to be restricted to conventional DC, as it is not found on plasmacytoid DC (Caminschi et al., 2008; Huysamen et al., 2008; Sancho et al., 2008). Thus, in the human, expression of XCR1 and Clec9A/DNGR-1 is not fully correlated, as is the case with conventional DC in the mouse. CADM1, another surface molecule tightly associated with Batf3-dependent cross-presenting DC in the mouse, gives a bright signal in the human and is perfectly correlated with CD141 (Galibert et al., 2005, and own unpublished data). Thus, it is possible that cross-presenting DC in the human can be demarcated by the co-expression of CD141, Clec9A, and CADM1. This assumption is further supported by the observation that SIRPα on DC in various human organs is correlated with CD1c and fully anti-correlated with CD141 (own unpublished data). Thus, on a broad scale, it is quite clear that the human CD141^+^ DC contain the cross-presenting DC population. Whether all of CD141^+^ DC can cross-present or whether this function is restricted to the XCR1^+^ CD141^+^ DC remains to be determined. Further detailed insight into this question will require gene expression and functional studies comparing the majority of CD141^+^ DC expressing XCR1 and the 20% fraction of CD141^+^ DC negative for XCR1.

## Conclusions and perspectives

In summary, recent developments in the field allowed major progress in the classification of mouse and human DC. Particularly in the mouse, where the subdivision of DC was notoriously difficult, the use of a general DC classification scheme based on the expression of XCR1 and SIRPα will make experimental results more precise and also more comparable. Without any doubt, the use of additional surface molecules will continue to be necessary in order to better understand the functional states of DC of a given lineage. Thus the old “markers” such as CD4, CD8, CD11b, or CD103 will not become obsolete, but they will obtain a new role in the functional analyses of XCR1^+^ or SIRPα^+^ mouse DC.

## Acknowledgement

The work was supported by grants of the Wilhelm Sander Foundation, the Fritz Thyssen Foundation, the VIP program of the Bundesministerium für Bildung und Forschung, and in part by the Deutsche Forschungsgemeinschaft (Kr 827/16-1 and Kr 827/18-1).

